# Pulmonary Fibrosis Ferret Model Demonstrates Sustained Fibrosis, Restrictive Physiology, and Aberrant Repair

**DOI:** 10.1101/2024.06.04.597198

**Authors:** Jacelyn E. Peabody Lever, Qian Li, Nikoleta Pavelkova, Shah S. Hussain, Sayan Bakshi, Janna Q. Ren, Luke I. Jones, Jared Kennemur, Mason Weupe, Javier Campos-Gomez, Liping Tang, Jeremie M.P. Lever, Dezhi Wang, Denise D. Stanford, Jeremy Foote, Kevin S. Harrod, Harrison Kim, Scott E. Phillips, Steven M. Rowe

## Abstract

**Rationale:** The role of MUC5B mucin expression in IPF pathogenesis is unknown. Bleomycin-exposed rodent models do not exhibit sustained fibrosis or airway remodeling. Unlike mice, ferrets have human-like distribution of MUC5B expressing cell types and natively express the risk-conferring variant that induces high MUC5B expression in humans. We hypothesized that ferrets would consequently exhibit aberrant repair to propagate fibrosis similar to human IPF.

**Methods:** Bleomycin (5U/kg) or saline-control was micro-sprayed intratracheally then wild-type ferrets were evaluated through 22 wks. Clinical phenotype was assessed with lung function. Fibrosis was assessed with µCT imaging and comparative histology with Ashcroft scoring. Airway remodeling was assessed with histology and quantitative immunofluorescence.

**Results:** Bleomycin ferrets exhibited sustained restrictive physiology including decreased inspiratory capacity, decreased compliance, and shifted Pressure-Volume loops through 22 wks. Volumetric µCT analysis revealed increased opacification of the lung bleomycin-ferrets. Histology showed extensive fibrotic injury that matured over time and MUC5B-positive cystic structures in the distal lung suggestive of honeycombing. Bleomycin ferrets had increased proportion of small airways that were double-positive for CCSP and alpha-tubulin compared to controls, indicating an aberrant ‘proximalization’ repair phenotype. Notably, this aberrant repair was associated with extent of fibrotic injury at the airway level.

**Conclusions:** Bleomycin-exposed ferrets exhibit sustained fibrosis through 22 wks and have pathologic features of IPF not found in rodents. Ferrets exhibited proximalization of the distal airways and other pathologic features characteristic of human IPF. MUC5B expression through native cell types may play a key role in promoting airway remodeling and lung injury in IPF.

## Introduction

Idiopathic Pulmonary Fibrosis (IPF) is a chronic, progressive interstitial lung disease of older adults. Disease progression is insidious; thus, most patients present late in their clinical course after the fibroproliferative process has caused extensive and permanent lung parenchymal damage. The mean survival is only 3-5 years and patients die of respiratory failure (1). Disease prevalence exceeds five million globally, and the incidence is increasing. The most common risk factor for developing IPF is a gain-of-function *MUC5B* promoter variant rs35705950, accounting for at least 30% of the total risk for developing IPF (2) and confirmed in independent studies (3–7)). The risk-conferring TT-minor allele variant rs35705950 is associated with enhanced *MUC5B* mucus expression in both unaffected subjects (2) and patients with IPF (8). Murine studies also suggest worsened fibrosis with ectopic Muc5b overexpression (9). However, an animal model with sustained fibrosis and human-like mucus distribution is unavailable to confirm these observations.

Much of our understanding about lung fibrosis originates from bleomycin (BLM) exposure in rodent animal models but the best characterized model (murine, single-dose of BLM administered oropharangeally) resolves spontaneously 3-4 weeks after injury and fails to recapitulate the hallmark lesions of IPF (10–12). Over 500 experimental studies describe the efficacy of novel compounds in the rodent-BLM model, yet therapies that reverse fibrotic lung damage and improve survival remain elusive, the modest effects of pirfenidone and nintedanib notwithstanding. Though no model system perfectly recapitulates complex human pathobiology, animal models that more closely mimic aberrant repair mechanisms of humans with IPF are needed to bridge translational gaps from the bench to the bedside (13).

We hypothesized that the mucus micro-environment was important for the propagation of sustained fibrosis and aberrant repair, but required an animal model where the physiology of mucus expression remained intact. Ferrets are mustelids and more evolutionarily similar to humans than rodents in their lung anatomy and epithelial cell biology. Relevant to the current study, ferrets have a similar distribution of MUC5B-producing submucosal glands as humans and express high amounts of MUC5B with native presence of the risk-conferring rs35705950 TT promoter variant (14).

Furthermore, ferrets provide advantages for modeling other airway diseases, including influenza (15), CF (16), and COPD (17), whereas rodent models do not recapitulate mucus-related defects. Thus, we sought to develop a ferret model of BLM–induced pulmonary fibrosis and measure clinically relevant parameters such as serial, longitudinal CT imaging and lung function analyses coupled with histopathological examination that examines epithelial cell metaplasia characteristic of aberrant repair in IPF. We found that ferrets exposed to a single dose of intra-tracheal (I.T.) BLM developed sustained fibrosis through 22 weeks and exhibit distal airway metaplasia with ‘proximalization’ of the distal airways and other pathologic features characteristic of human IPF that are absent in mice. These results affirm the importance of mucin expression in aberrant IPF remodeling, and provides BLM exposed ferrets as a research tool to address an unmet need in IPF by developing a more human-relevant model to study pathogenesis at the cellular level.

## Methods

### In Vivo Bronchoscopy and Intra-tracheal Delivery of Bleomycin Sulfate or Saline

Age- and sex-matched wild-type ferrets (*Mustela putorius furo*, body weight 600– 900g in females, 1200–2000g in males) were purchased from Marshall BioResources and randomly assigned to receive bleomycin (BLM) or saline control. To optimize delivery and control for sexually dimorphic ferrets, dosing was conducted in both weight and length specific manners. A custom-built miniaturized video-bronchoscopy system (PolyDiagnotics, Germany) with an outer diameter of 1.75 mm was used to measure the carina location, enabling length-specific dosing. A single dose of bleomycin sulfate (5U/kg), in sterile endotoxin-free saline (400 µl) or control sterile endotoxin-free saline (400 μl) was instilled with a microsprayer I.T. (Biojane, China) on anesthetized ferrets 2 cm from carina.

### Clinical Outcomes

Body weight (Ohaus) and resting pulse oximetry (Waveline Nano-V2, Avante, Kentucky, USA) were measured to assess clinical phenotype in ferrets longitudinally. Additionally, depth of anesthesia during procedures was monitored with pulse oximetry.

### Standardized Anesthesia and Intubation

Ferrets were anesthetized as previously described using a formulation of dexmedetomidine (0.08 – 0.2 mg/kg, IM) in combination with ketamine (2.5-5 mg/kg, IM) for studies requiring intubation or isoflurane by inhalation (2-3% with O_2_) for µCT (17, 18). Ophthalmic lubricant was applied to the eyes once sedation was achieved, and the animals were placed on a heating pad until recovery. Anesthetic reversal was achieved with atipamezole (5.0 mg/ml), dosed at an equal volume as the dexmedetomidine and supplemental oxygen was provided as necessary. Ferrets were intubated with a 2.5-mm uncuffed endotracheal tube for treatment installation or with a 3.0-mm cuffed endotracheal tube (Medtronic Shiley, Minnesota, USA) for flexiVent using a laryngoscope equipped with a size 0 Miller blade (Welch Allyn). Endotracheal intubation was confirmed with a pediatric end-tidal CO_2_ detector (Medtronic, Minnesota, USA).

### Lung Function Measurements

Lung function in ferrets was characterized using the flexiVent mechanical ventilator (SCIREQ), as previously described (17). Briefly, ferrets were sedated, intubated with 3.0-mm cuffed endotracheal tubes, and connected to the in-line ventilator for multiple forced oscillatory maneuvers to quantify pressure-volume loops, inspiratory capacity, resistance, compliance, and elastance. Ferrets remained on ventilator support until they recovered from sedation and were able to breathe spontaneously. Animals had serial measurements conducted longitudinally at baseline and through 22-weeks post treatment.

### CT Imaging

We conducted non-contrast, respiratory-gated CT imaging with ultra-focused magnification, on spontaneously breathing ferrets anesthetized under inhaled isoflurane (MiLabs, Utrecht, The Netherlands), as previously described (18). All images were acquired with the following parameters: tube voltage (55 kV), tube current (0.19 mA), scan angle (360°), and 20-30 ms of exposure. All images were reconstructed with MiLabs software at 80 μm/voxel at the single respiratory phase corresponding with maximal inspiration. Post reconstruction, lung masks were generated in the PMOD software (View and 3D modules; PMOD Technologies LLC, Zurich, Switzerland) for quantitative volumetric analysis of lung opacification by Hounsfield units (HU). The data is displayed using the standard human lung window (Window Width: 1600, Window Level: −600, HU range: −1350-150).

### Euthanasia and Necropsy

Ferrets were euthanized with injectable anesthetics (dexmedetomidine and ketamine) followed by bilateral thoracotomy and exsanguination, conforming to the guidelines of the *AVMA Panel on Euthanasia*. Separate lung lobes were designated for histopathology and biochemical assays. Lungs were inflated, fixed, and embedded as previously described (17).

### Histopathology and Microscopy

Hematoxylin and eosin (H&E) and Masson’s trichrome staining were performed on paraffin-embedded lung sections. A board-certified veterinary pathologist assessed histopathologic changes (see supplemental methods for detailed criteria) and quantitative morphometry were assessed by blinded reviewers using the modified Ashcroft score (19). Images were obtained from Lionheart FX (Biotek) in the UAB High Resolution Imaging Facility and the UAB Cystic Fibrosis Research Center.

### Immunofluorescent Staining and Analysis

Airway remodeling was examined by co-localization of alpha-tubulin (AT), a marker of ciliated cells found in medium and large airways, and club cell secretory protein (CCSP), a marker for non-ciliated bronchiolar epithelium of small airways, by dual immunofluorescence. Unstained paraffin-embedded lung slices were fluorescently labeled using standard IF process with antigen retrieval (see supplemental methods for detailed process and reagent list).

For the semi-automated analysis, of the percentage of AT or CCSP signal along the apical surface of the airway epithelia (AT+%, CCSP+% or AT+,CCSP+%) were estimated using Image J (NIH) and scripts written in NI LabVIEW^TM^ 2024 Q1 (Emerson). See supplemental methods for details and overview of manual check points for quality control.

For validation/quality control, we manually quantified the proportion of airways, controlling for diameter, that were single positive for either AT and CCSP and those that were AT+CCSP+ double positive and recorded the modified Ashcroft Score associated with each airway in a subset of ferrets.

### Statistics

Descriptive statistics (mean, SD, and SEM) were compared using Two-Way ANOVA or mixed-effects model with the Geisser-Greenhouse correction, as appropriate. Post-hoc tests for multiple comparisons were calculated using Dunnett’s multiple comparison’s test when the ANOVA was significant in GraphPad Prism. All statistical tests were 2-sided and were performed at a 5% significance level. Error bars designate SEM unless indicated noted.

### Study Approval

All animal protocols used in this work were reviewed and approved by the University of Alabama at Birmingham Institutional Animal Care and Use Committee.

## Results

### Clinical Phenotype

We developed a ferret model of pulmonary fibrosis by delivering a single dose of BLM, in a weight and length specific (5U/kg, 2 cm from carina). Pulse oxygen saturation and weights were measured longitudinally in control and bleomycin animals. Data were analyzed with mixed-effects model with Geisser-Greenhouse correction and post-hoc analysis was performed with Sidak’s multiple comparison test to compare control and bleomycin ferrets at each timepoint. For pulse oximetry, we found a significant fixed effect of timepoint (P<0.0001), treatment group (P<0.001), and interaction between timepoint and treatment group (P=0.0003). There was no difference between control and bleomycin at baseline by Sidak post-hoc (mean difference [95%CI of difference]: 0.12%[-0.1263-0.3570], P=0.7066). However, at 3, 6, 12, 18, and 22 weeks following treatment with saline or BLM, the BLM group had statistically significantly lower oxygen saturation compared to contemporaneous controls (mean difference [95%CI of difference]- 3wk: 6.94%[4.056-9.814], P<0.0001; 6wk: 4.54%[2.405-9.814], P<0.0001; 12wk: 4.62% [1.030-8.199], P=0.0076; 18wk 9.13% [2.477-15.79], P=0.0125; 22wk: 10.20% [0.1551-20.24], P=0.0474) (**Figure 1A**).

**Figure 1.**
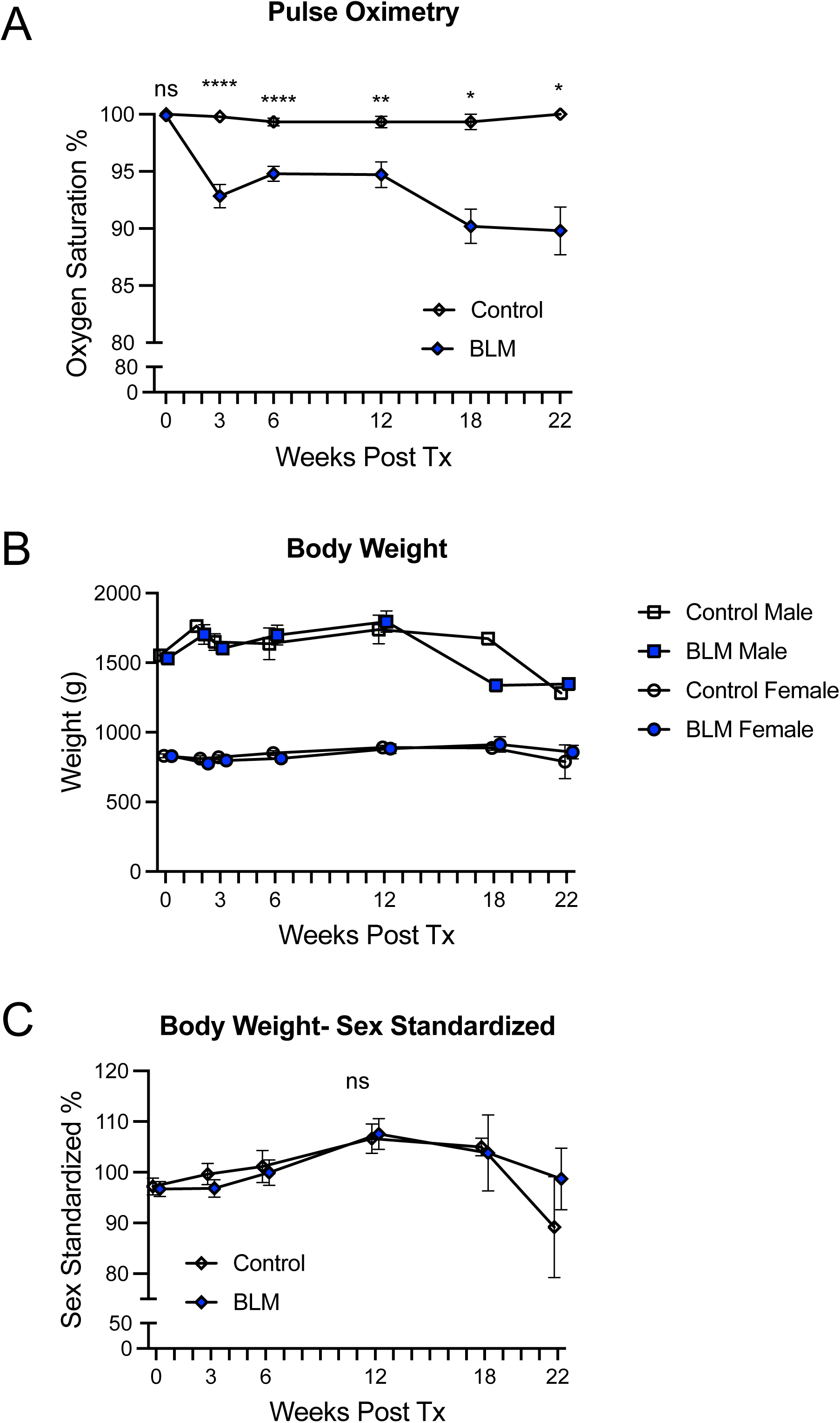
The clinical phenotype of BLM-PF ferrets is decreased blood oxygenation by pulse oximetry, but body weight is unaffected. **A)** Pulse oximetry in saline control and bleomycin (BLM) ferrets was measured at baseline, 3-weeks, 6-weeks, 12-weeks, 18-weeks, and 22-weeks (N=27 control; N=5-35 BLM ferrets per timepoint). Significant decrements in blood oxygenation were observed in the BLM ferrets 3, 6, 12, 18, and 22-weeks post treatment relative to control; ****P<0.0001, ***P<0.001, **P<0.01, *P<0.05, ns=not significant. Oxygen saturation did not differ between control and BLM groups at baseline. **B)** Sex split, raw weights are shown for control and BLM ferrets over time. Ferrets are sexually dimorphic, but there was no difference between treatment groups. **C)** To allow for male and female data to be pooled by group, body weight was sex standardized by the sex-specific average weight of baseline BLM and control animals (repeated measures averaged for control). There was no significant difference by treatment group or significant interaction between timepoint and treatment group.

Due to the sexual dimorphism of ferrets, raw body weight over time by treatment group is split by sex (**Figure 1B**). To allow for male and female weight data to be pooled by group for statistical analysis, body weight was sex standardized by the sex-specific average weight of baseline BLM and control animals, with repeated measures averaged for control. There was a significant fixed effect of timepoint (P<0.0001) but treatment group and interaction between timepoint and treatment group were not statistically significantly different by mixed-effects model with Geisser-Greenhouse correction (P=0.8116 and P=0.5174, respectively).There was no difference between control and bleomycin sex standardized weights at baseline, 3, 6, 12, 18, or 22 weeks following their respective treatments by Sidak (mean difference [95%CI of difference] (mean difference [95%CI of difference]-Baseline: 0.492%[-0.5.431 to −6.414], P>0.9999); 3wk: 2.821%[-4.511 to −10.15], P=0.8837; 6wk: 1.207%[-9.882 to −12.3], P<0.998; 12wk: - 0.925% [-12.57to −10.72], P>0.9999; 18wk: 1.186% [-33.56 to −35.93], P>0.9999; 22wk: −9.526% [-71.41 to −52.36], P=0.9766) (**Figure 1C**).

### Sustained Restrictive Physiology in BLM-Ferrets

Considering that IPF patients demonstrate a restrictive pattern on pulmonary function tests, including decreased lung volumes, reduced lung dynamic and static compliance, increased elastance and static recoil pressure (20), we decided to investigate pulmonary pathophysiology over the experimental time-course. We performed various forced oscillation techniques in ferrets intubated with cuffed endotracheal tubes (**Figure 2A**). We assessed inspiratory capacity (IC), by inflating the lungs to a total lung capacity state (a pressure of 30 cmH_2_O), allowed alveolar pressure to equilibrate, and recorded the difference between initial and end volumes. Since, ferrets are sexually dimorphic and lung volumes are influenced by lung size (21), we display male and female raw values separately (**Figure 2B**). To allow for male and female IC data to be pooled by group for statistical analysis, IC volume was sex standardized by the sex-specific average IC of baseline BLM and control animals, with repeated measures averaged for control and data are displayed as a sex standardized percent of expected. Data were analyzed with mixed-effects model (P<0.0001) and post-hoc analysis was performed with Dunnett’s multiple comparison test to compare baseline BLM IC with subsequent BLM timepoints and control. There was no difference between baseline BLM and control IC sex standardized % (mean [95%CI]: Baseline BLM 97.73% [90.29-105.2]; Control 101.1% [96.76-105.4]; mean difference [95%CI] 3.44 [-7.194-14.08], P=0.9334). However, there was significant, sustained decrease in IC sex standardized % of expected at 3, 6, 12, 18, and 22 weeks post-BLM compared to BLM baseline (mean difference [95%CI of difference]- 3wk: −21.66%[-28.93 to −14.4], P<0.0001; 6wk: −10.89%[-18.29 to −3.485], P=0.0012; 12wk: −14.4% [-22.52 to −6.273], P<0.0001; 18wk −15.25% [-27.10 to −3.397], P=0.0057; 22wk: −17.93% [-29.78 to - 6.075], P=0.0008) (**Figure 2C**).

**Figure 2.**
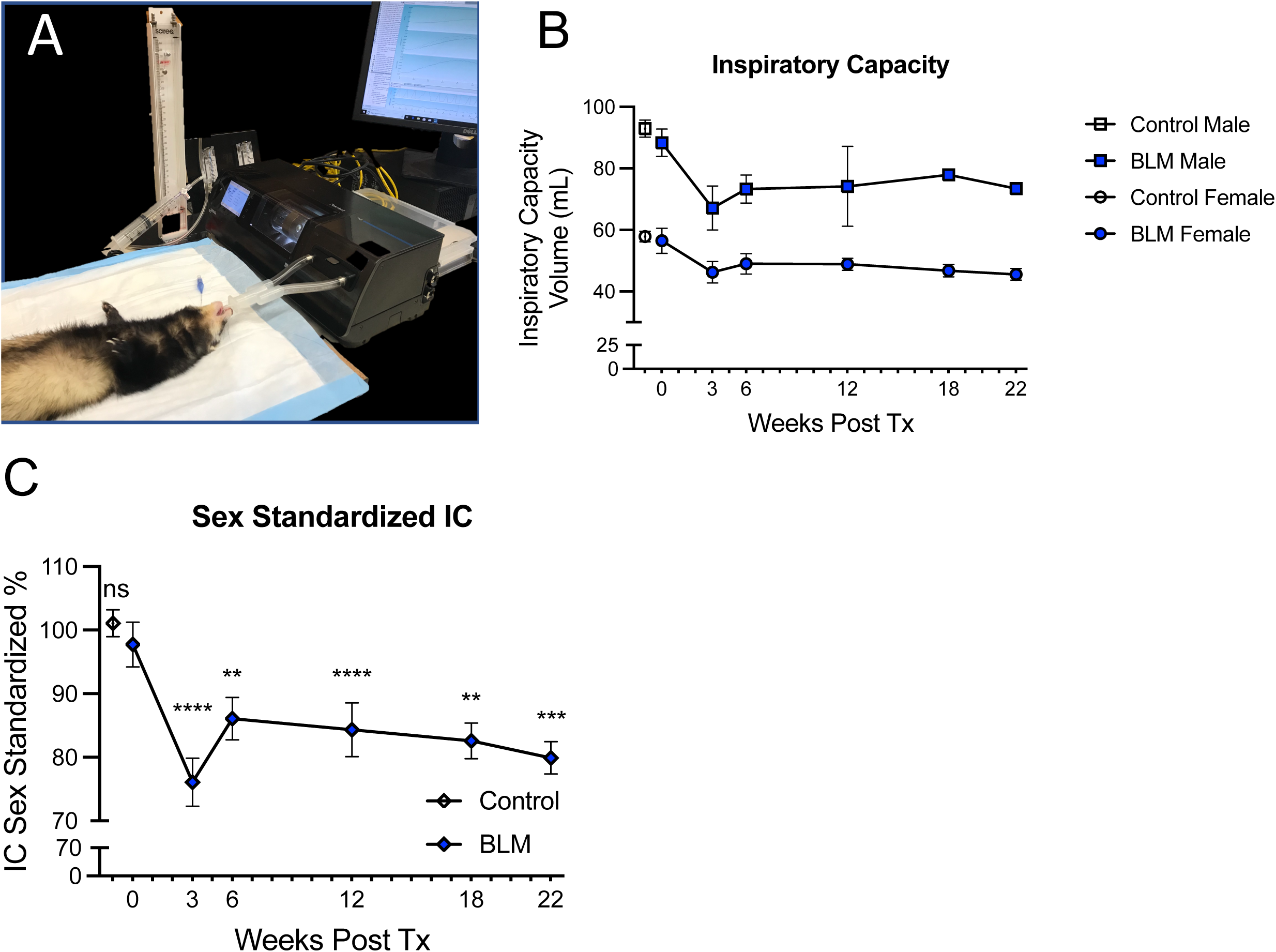
BLM-ferrets have sustained inspiratory capacity decrement. **A)** Photograph of experimental setup for flexiVent lung function measurements. The ferret is intubated with a 3.0mm cuffed endotracheal tube and mechanically ventilated, then volume and pressure measurements are recorded. For inspiratory capacity (IC), a deep inflation maneuver is used to gradually inflate the lungs to a total lung capacity state (defined as pressure of 30 cmH_2_O), followed by a hold to allow equilibration of alveolar pressure. IC is calculated using the measured initial and end volumes. **B)** Sex split, raw IC volumes are shown for control and BLM ferrets over time. Graphical representation Male ferrets represented graphically with a square, female ferrets with a circle**. C)** To allow for male and female data to be pooled by group, IC volumes were sex standardized by the sex-specific average IC of baseline BLM and control animals (repeated measures averaged for control). IC was measured at 3-weeks, 6-weeks, 12-weeks, 18-weeks, and 22-weeks (N=27 control ferrets, repeated measures averaged; N=5-18 BLM ferrets per timepoint. Control and BLM baseline IC sex standardized % was not different. However, there was significant decrease in IC sex standardized % of expected at 3, 6-, 12-, 18-, and 22-weeks post-BLM relative to BLM baseline; ****P<0.0001, ***P<0.001, **P<0.01, *P<0.05, ns=not significant. Sex pooled data represented graphically with a diamond.

Pressure Volume (PV) loops were measured serially in saline control and BLM-treated ferrets at baseline, 3-weeks, 18-weeks, and 22-weeks using combined respiratory physiology via the flexiVent system (see **Figure 2A** for experimental setup). PV loops were measured by delivering a series of pressure-driven, stepwise maneuvers. Briefly, the baseline pressure was set to PEEP (to model functional residual capacity), lungs were inflated over 8 seconds in a stepwise fashion until pressure reached 30 cmH2O (total lung capacity was reached), and then the lungs are deflated in a similar stepwise fashion over 8 seconds until pressure was back to baseline. See **Figure 3A** for representative serial pressure volume (PV) loops from an individual control female ferret and an individual BLM female ferret at baseline (D0), 3-weeks, 18-weeks, and 22-weeks post treatment.

**Figure 3.**
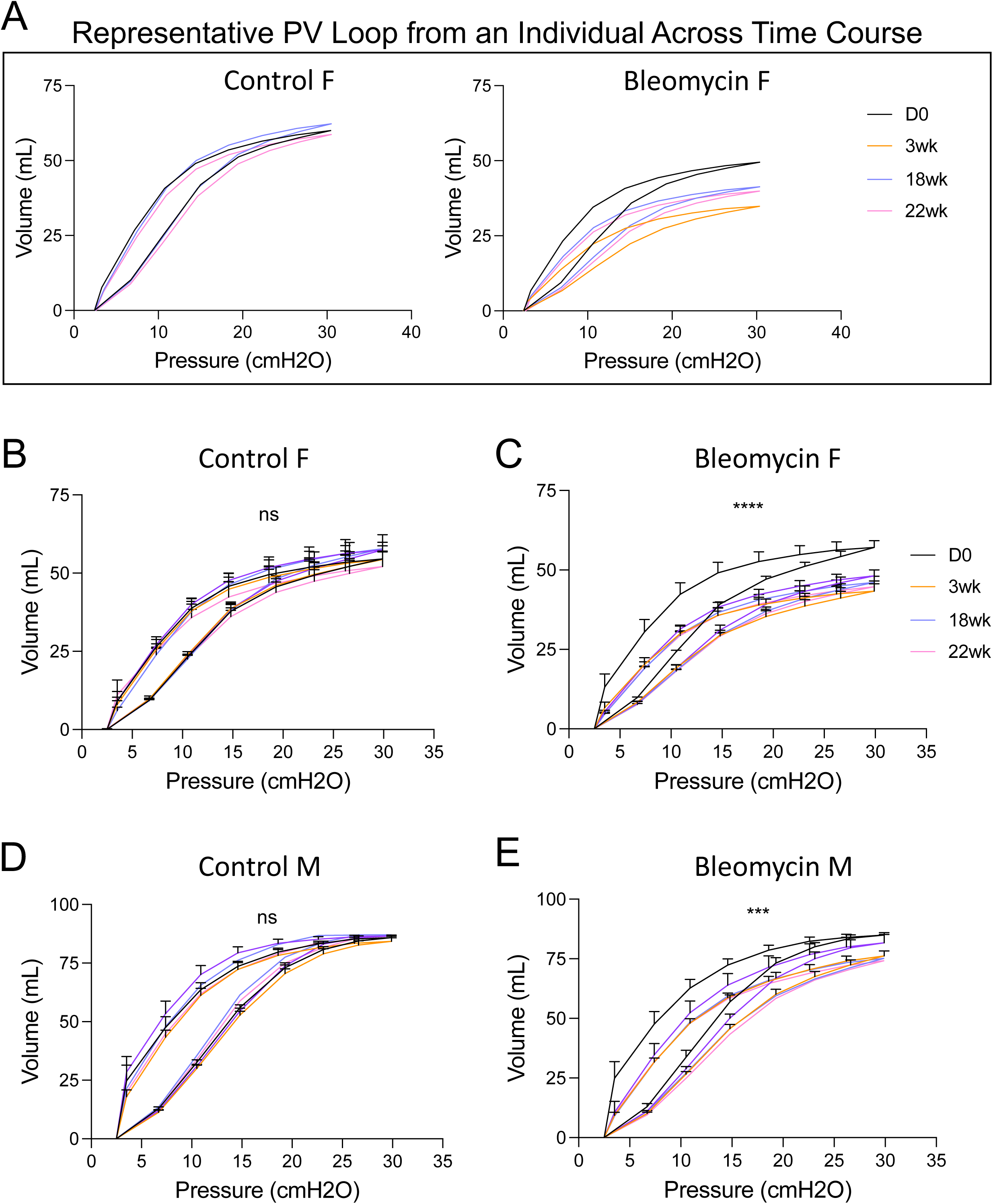
Pressure-Volume loop of BLM-ferrets demonstrate sustained restrictive physiologic changes. Using the same experimental setup for flexiVent lung function measurements shown in Figure 2A, we derived Pressure Volume (PV) loop by delivering a series of pressure-driven, stepwise maneuvers. The baseline pressure is the functional residual capacity (matched to PEEP), lungs are inflated over 8 seconds in a stepwise fashion until pressure of 30 cmH_2_O (total lung capacity), and then the lungs are deflated in a similar stepwise fashion over 8 seconds until baseline pressure is reached. **A)** Representative PV curves from an individual control female ferret and an individual BLM female ferret at baseline (D0), 3-weeks, 18-weeks, and 22-weeks post treatment. Note the reproducibility in the serial PV loops in the control animal and the persistent downward shift of the BLM curves characterized by decreased volumes at the same pressure at 3, 18, and 22 weeks compared to her baseline. **B)** Pooled PV loops for control female ferrets at baseline (N=9), 3-weeks (N=8), 18-weeks (N=4), and 22-weeks (N=4); P=0.8150, ns. **C)** Pooled PV loops for bleomycin female ferrets at baseline (N=12), 3-weeks (N=12), 18-weeks (N=4), and 22-weeks (N=4); ****P<0.0001. **D)** Pooled PV loops for control male ferrets at baseline (N=11), 3-weeks (N=13), 18-weeks (N=1), and 22-weeks (N=1); P=0.0778, ns. **E)** Pooled PV loops for bleomycin male ferrets at baseline (N=13), 3-weeks (N=14), 18-weeks (N=1), and 22-weeks (N=1); ***P<0.0007.

PV loop data were analyzed longitudinally by treatment group and sex using a mixed-effects model with Geisser-Greenhouse correction. There was no significant difference in PV loops between timepoints for control females (P=0.8150) **(Figure 3B)** or control males over time (P=0.0778) (**Figure 3D)**. In comparison, there is a downward shift of the PV loops following BLM treatment compared to baseline in both the BLM females (P<0.0001) (**Figure 3C)** and BLM males (P=0.0007) (**Figure 3E)**, indicative of restrictive change for which greater pressure is required to inflate the lungs to the same volume.

We also performed sinusoidal wave, single-frequency oscillation maneuvers with flexiVent. The volume of gas delivered to the ferret and the flow of gas at the airway opening is experimentally controlled and pressure at the airway opening is recorded. From this, we can determine dynamic respiratory system compliance (Crs), elastance (Ers), and resistance (Rrs) using the single-compartment linear model (Pressure_Trachea_= (Resistance*Flow_trachea_) + (Elastance*Volume_tracheal_) + PEEP) (22, 23).

Compliance (Crs) is defined by the change in volume that occurs per unit pressure and describes the ease of respiratory distension (Compliance=1/elastance). Sexual dimorphism is apparent when comparing raw Crs values for control and BLM ferrets, split by sex (**Figure 4A)**. To allow for male and female Crs data to be pooled by treatment group for statistical analysis, Crs (mL/cmH_2_O) was sex standardized in same manner as was conducted for IC. Data were analyzed with mixed-effects model (P<0.0001) and post-hoc analysis was performed with Dunnett’s multiple comparison test to compare baseline BLM Crs with subsequent BLM timepoints and control. There was no difference between baseline BLM and control Crs sex standardized % (mean difference [95%CI] 1.843[-7.398 to 11.08], P=0.9958). However, there was significant decrease in Crs sex standardized % of expected at 3, 6, 12, 18, and 22 weeks post-BLM compared to BLM baseline (mean difference [95%CI of difference]- 3wk: −15.27%[-20.08 to −10.47], P<0.0001; 6wk: −13.38%[-18.43 to −8.333], P<0.0001; 12wk: −12.24% [-17.70 to −6.785], P<0.0001; 18wk −15.10% [-24.35 to −5.846], P=0.0002; 22wk: −14.38% [-23.63 to −5.125], P=0.0005) (**Figure 4B**).

**Figure 4.**
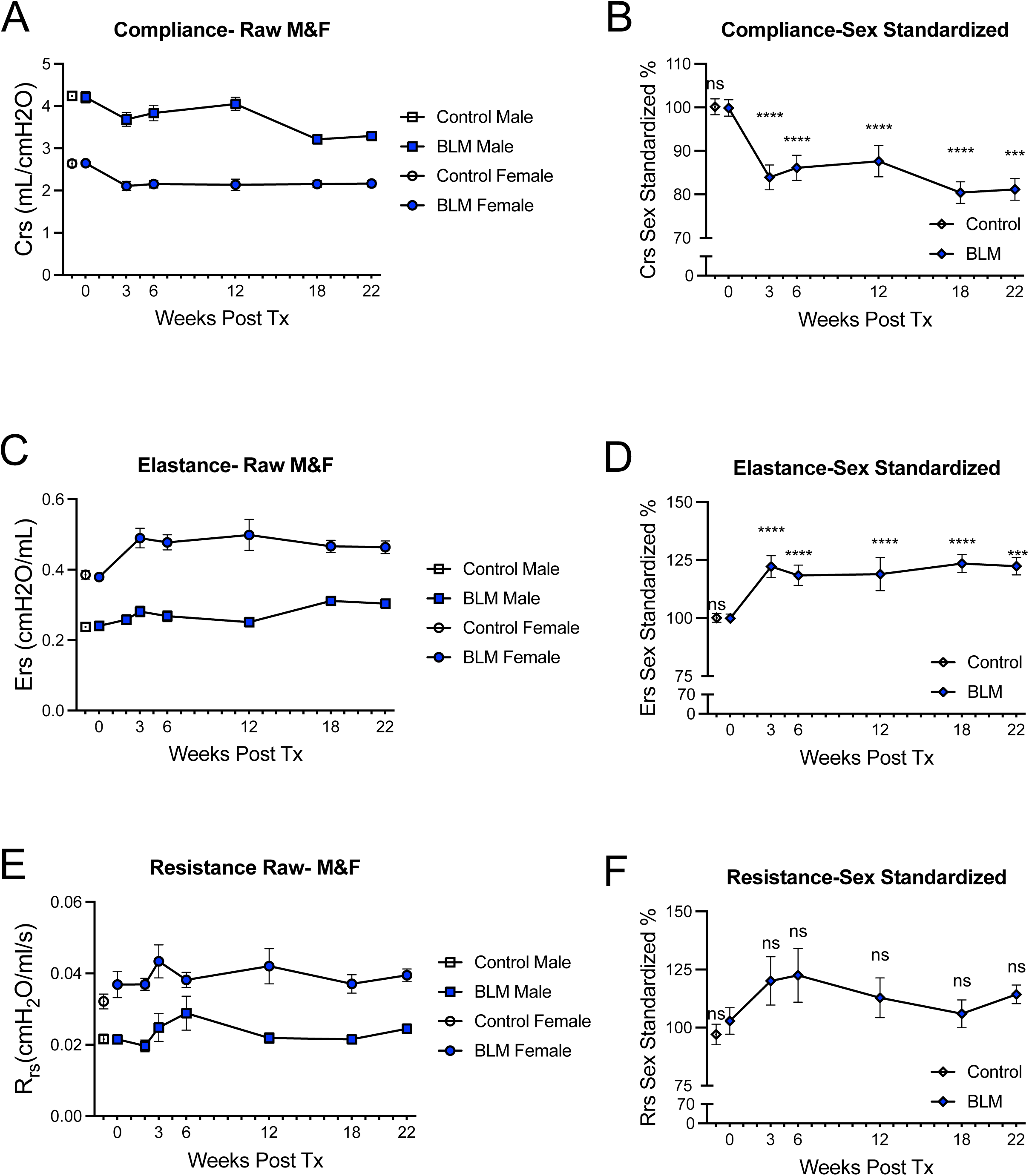
BLM-ferrets recapitulate restrictive respiratory mechanics, characterized by decreased dynamic lung compliance and increased elastance, with no change in total respiratory system resistance by single-compartment model *in vivo*. Sinusoidal wave, single-frequency oscillation maneuvers were conducted via flexiVent to determine dynamic respiratory system compliance (Crs), elastance (Ers), and resistance (Rrs) using the single-compartment linear model. Sex split, raw parameters are shown for control and BLM ferrets over time for **A)**Crs, **C)** Ers, and **E)** Rrs. Graphical representation Male ferrets represented graphically with a square, female ferrets with a circle**. B,D,F)** Crs, Ers, and Rrs were measured at 3-weeks, 6-weeks, 12-weeks, 18-weeks, and 22-weeks (N=27 control ferrets, repeated measures averaged; N=5-18 BLM ferrets per timepoint. Male and female data were pooled by group, following sex standardization by the sex-specific parameter average of baseline BLM and control animals (repeated measures averaged for control). **B)** Control and BLM baseline Crs sex standardized % was not different. However, there was significant decrease in Crs sex standardized % of expected at 3, 6-, 12-, 18-, and 22-weeks post-BLM relative to BLM baseline; ****P<0.0001, ***P<0.0005, ns=not significant. **D)** Control and BLM baseline Ers sex standardized % was not different. However, there was significant increase in Ers sex standardized % of expected at 3, 6-, 12-, 18-, and 22-weeks post-BLM relative to BLM baseline; ****P<0.0001, ***P<0.0002, ns=not significant. **F)** Control and BLM baseline Rrs sex standardized % was not different and there was no difference between BLM baseline and 3, 6-, 12-, 18-, or 22-weeks post-BLM relative to BLM baseline; ns=not significant.

Elastance (Ers) is inversely proportional to compliance and increases in interstitial lung disease due to the pathopneumonic deposition of extracellular matrix which characterizes these diseases. Raw Ers values for control and BLM ferrets, split by sex are shown in **Figure 4C**. The raw Ers (cmH_2_O/mL) data were subsequently sex standardized and pooled by treatment group. Data were analyzed with mixed-effects model (P<0.0001) and post-hoc analysis was performed with Dunnett’s multiple comparison test to compare baseline BLM Ers with subsequent BLM timepoints and control. There was no difference between baseline BLM and control Ers sex standardized % (mean difference [95%CI] 3.165 [-13.96 to 20.29], P=0.9972). However, there was significant increase in Ers sex standardized % of expected at 3, 6, 12, 18, and 22 weeks post-BLM compared to BLM baseline (mean difference [95%CI of difference]- 3wk: −28.91%[-37.52 to −20.30], P<0.0001; 6wk: −25.57%[-34.43 to −16.72], P<0.0001; 12wk: −25.78% [-35.48 to −9.265], P<0.0001; 18wk −22.38% [-35.50 to - 9.265], P=0.0001; 22wk: −21.71% [-34.83 to −8.590], P=0.0002) (**Figure 4D**).

Resistance (Rrs) depends on airway length and radius thus it is a marker of airway constriction and is influenced by sexual dimorphism. Raw Rrs values for control and BLM ferrets, split by sex are shown in **Figure 4E**. The raw Rrs (cmH_2_O/mL/s) data were subsequently sex standardized and pooled by treatment group. Data were analyzed with mixed-effects model (P=0.1732) and post-hoc analysis was performed with Dunnett’s multiple comparison test to compare baseline BLM Rrs with subsequent BLM timepoints and control. There was no difference between baseline BLM and control Rrs sex standardized % (mean difference [95%CI] −6.642 [-36.23 to 22.95], P=0.9972). Additionally and as expected for the nature of BLM induced injury, there were no significant differences in Rrs sex standardized % of expected at 3, 6, 12, 18, and 22 weeks post-BLM compared to BLM baseline (mean difference [95%CI of difference]- 3wk: 16.5%[-10.68 to 43.69], P=0.4621; 6wk: 20.09%[-8.601 to 48.78], P=0.3044; 12wk: 11.90% [-17.87 to −41.67], P<0.8501; 18wk −8.185% [-58.02 to 41.65], P=0.9987; 22wk: −21.71% [-49.67 to 50.00], P>0.9999) (**Figure 4F**).

### Evidence of Sustained Fibrosis in BLM-exposed ferrets

We conducted longitudinal, respiratory-gated µCT scans to visualize pulmonary fibrosis development *in vivo*. We observed heterogenous fibrotic change, characterized by airway-centric, reticular opacities with patchy areas of ground-glass opacities with some instances of possible subpleural honeycomb change, persistent through 22- weeks (**Figure 5A)**. We performed volumetric µCT analysis and calculated the 85^th^ percentile density index (P85) within the lung masks to quantify this opacification (**Figure 5B** demonstrates P85 on HU histogram for a representative animal over time). Data were analyzed with One-way ANOVA (P<0.0001) and post-hoc analysis was performed with Sidak’s multiple comparison test to compare baseline BLM P85 HU with subsequent BLM timepoints and control. Relative to BLM baseline, P85 increased by 17.07% at 3wks-post-BLM (mean difference [95%CI]; 90.63 HU [15.15 to 166.1], P=0.0114), 37.66% at 12wks-post-BLM (mean difference [95%CI]; 199.9 HU [119.4 to 280.4], P<0.0001), 44.65% at 18wks-post-BLM (mean difference [95%CI]; 237.0 HU [127.9 to 346.2], P<0.0001), and 38.02% at 22wks-post-BLM (mean difference [95%CI]; 201.8HU [92.63 to 310.9], P<0.0001) (**Figure 5C)**. Control ferrets were not statistically different from BLM baseline (mean difference [95%CI]; 8.03 HU [-68.91 to 84.97], P=0.9995).

**Figure 5.**
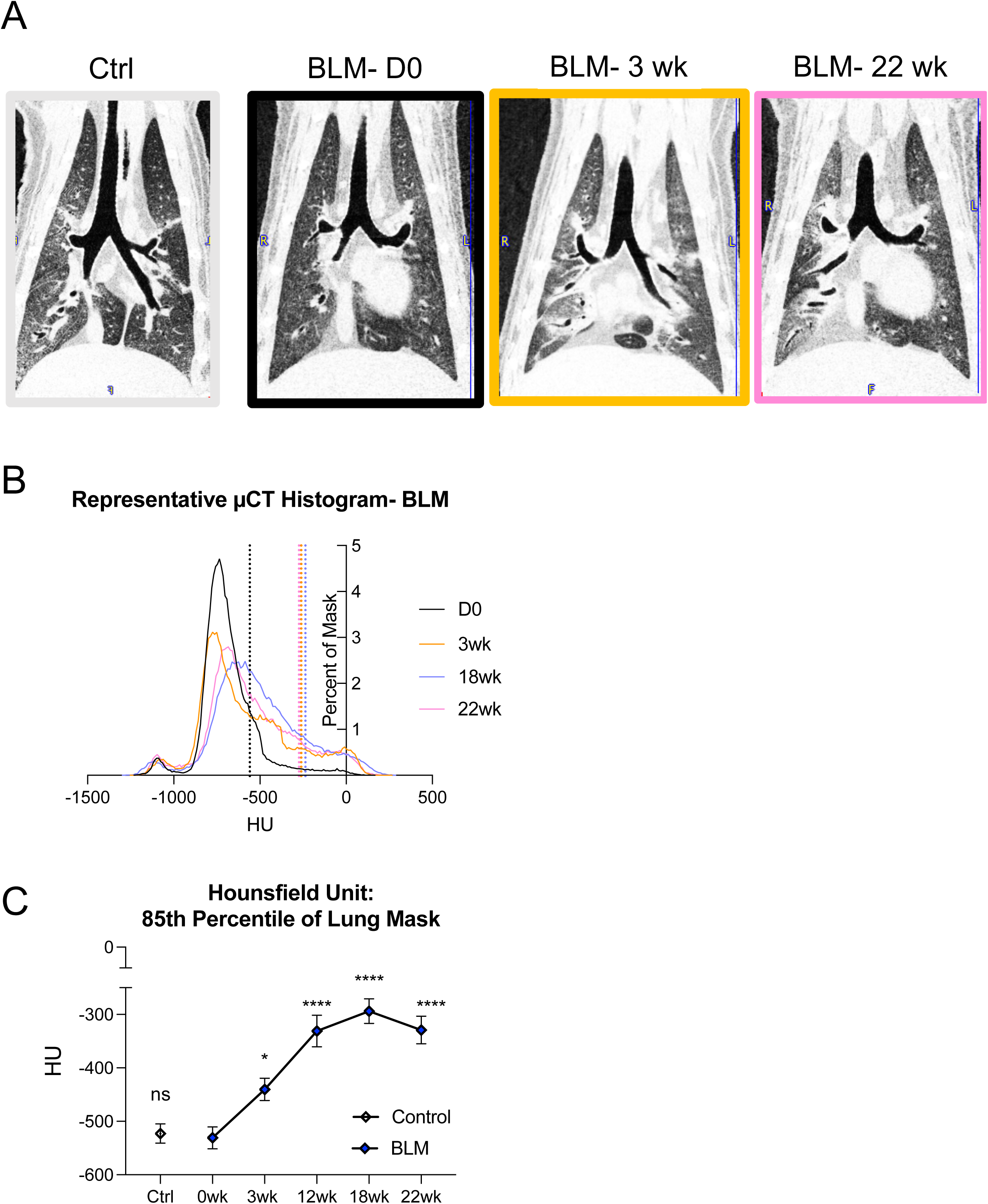
Radiographic opacification of the lungs is persistent through 22-weeks in BLM-ferrets by quantitative, volumetric analysis of respiratory-gated µCT scans. **A)** Representative 2D slices of a control ferret and serial scans of BLM ferret over time. **B)** Representative histogram time course of a BLM ferret generated from the 3D lung mask; dotted line denotes the 85th percentile density index (P85). Note the sustained rightward shift of P85, indicative of increased opacification of the lung volume. **C)** Quantification of P85. Control P85 HU was not different than BLM baseline. However, there was significant increase in P85 HU at 3-, 12-, 18-, and 22-weeks post-BLM relative to BLM baseline; ****P<0.0001, *P=0.0114, ns=not significant.

To further bolster these *in vivo* findings of longitudinal human-relevant imaging evidence of fibrosis, we additionally evaluated lungs grossly (**Figure 6A-6D**), histologically (**Supplemental Figure 1A,B)** including by Masson’s Trichrome **(Figure 6E-6H)**, biochemically by quantifying hydroxyproline content of lung homogenate (**Supplemental Figure 2A),** and with immunohistochemistry of hydroxyproline and immunofluorescent staining of alpha-smooth muscle actin, (**Supplemental Figure 2B-M.**

**Figure 6.**
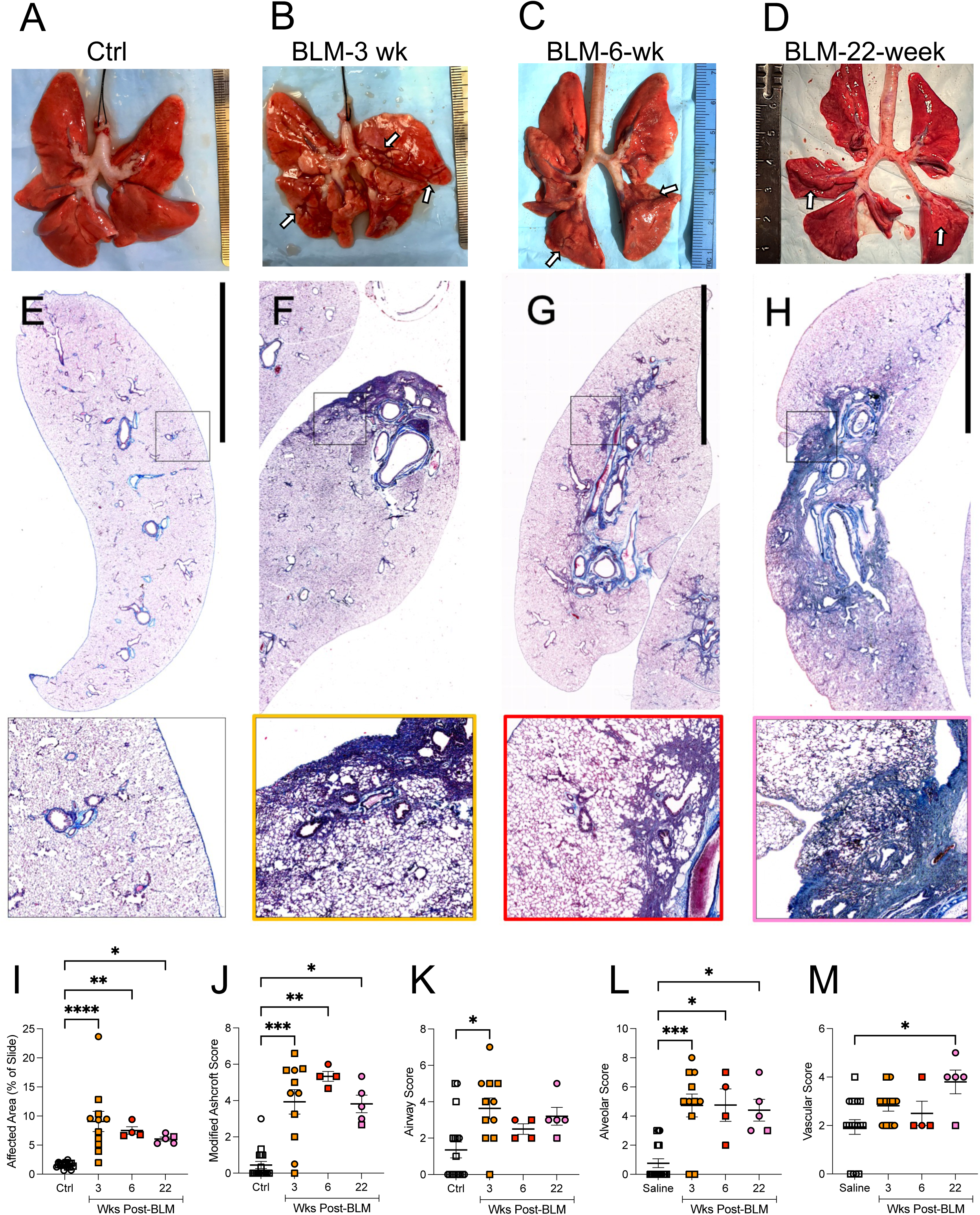
BLM-exposure induces fibrosis and collagen deposition in ferret lungs. Ferrets were exposed to a single dose of saline or 5U/kg BLM I.T. 2cm from carina via microspray and a lower lobe was allocated for histologic examination at 3, 6 and 22wk (N =11, 4 and 5, respectively, and 18 for saline group) as described in Methods. **A-D)** Representative gross prosection of whole lungs after BLM exposure shows subpleural fibrotic lobulation at 3 (**B**), 6 (**C**), and 22 weeks (**D**) (white arrows) compared to saline controls (**A**) (scale ruler with each photo). E-H) Typical photomicrographs of Masson’s trichrome staining of lungs from saline (E)- and BLM-treated ferrets at 3 (**F**), 6 (**G**) and 22 weeks (**H**), 4X magnification (scale bar 1cm), with square inlay enlarged. Semi-quantitative analysis of histology was perform by blinded, board certified veterinary pathologist for % fibrotic area (**I**) and Modified Ashcroft score (J) from Masson’s trichrome, and scores for airway (**K**), alveoli (**L**) and vasculature **M**) sublocations from H&E staining. ****P <0.0001 for (**I**) and (**J**); = 0.0059, 0.0002, and 0.0318 for (**K**), (**L**), and (E), respectively, by one-way ANOVA (Kruskal-Wallis test). *P<0.05; **P<0.01; ***P<0.001; ****P<0.0001 by Dunn’s test. Male ferrets represented by squares and female ferrets represented by circles.

Grossly, the pleural surface of the BLM lungs shows retraction along interlobular septa resulting in fibrotic lobulation appearance at 3, 6, and 22-weeks post BLM, compared to the smooth pleural surface of control (**Figure 6**). Of note, smaller, more cobblestone-like lesions were noted at 22-weeks, indicating more distorted lung architecture (**Figure 6D**; see also **Supplemental Figure 1B”** for the histological corollary of the interlobular septa retraction).

Histologically, BLM induced peribronchiolar and alveolar interstitial fibrosis, type II pneumocyte and bronchiolar epithelial hyperplasia, and edema as early as 3-weeks post-exposure, which matured to persistent fibrotic lesions through 22-weeks.

Additionally, the ferret model recapitulated pathology found in in human IPF including MUC5B+ AT+ mucociliary cystic structures in the distal lung reminiscent of IPF honeycomb cysts and fibroblastic foci (**Supplemental Figure 1B’**). At 3-weeks post BLM, alveolar septa showed fibrotic change with knot-like formation and some contiguous thickened regions, isolated fibrotic masses, inflammatory infiltrate resulting in hyper-cellular appearance, and collagen deposition (**Figure 6F**). At 6-weeks post BLM, there was less cellular infiltrate and collagen deposition, most prominent in dependent airways but extending to the pleura (**Figure 6G).** At 22-weeks post BLM, the lung architecture was severely damaged, alveoli were partly enlarged and rarefied, there were notable confluent fibrotic masses, and fibrotic banding connecting bronchioles lobular septa, and dense collagen deposition (**Figure 6H**).

Semi-quantitative histologic morphometric analysis conducted by a blinded, veterinary pathologist further supports the persistent nature of the lesions. Percent of fibrosis affected area for up to four lung slices increased from 1.6%±0.1 in control lungs to 9.1%±1.7, 7.5%±0.6, and 6.0%±0.3 in BLM lungs at 3, 6 and 22wk, respectively (**Figure 6I**, P<0.0001 by one-way ANOVA with non-parametric test; P<0.0001, =0.0075, and 0.0424 for 3, 6 and 22wk BLM lungs compared to saline lungs, respectively).

Consistently, Modified Ashcroft score for representative areas of the same lung slices also increased from 0.3±0.1 in saline lungs to 3.9±0.7, 5.3±0.3, and 4.5±0.3 in BLM lungs at 3, 6 and 22wk, respectively (**Figure 6J**, P<0.0001 by one-way ANOVA with non-parametric test; P=0.0011, 0.0022, and 0.0183 for 3, 6 and 22wk BLM lungs compared to saline lungs, respectively). Therefore, both affected area and severity of the fibrotic injury persisted up to 22wk post BLM exposure in ferrets.

Lesions were further subdivided into three anatomic sublocations (airway, alveoli, and vascular) and scored (**Figure 6K, 6L, 6M**). The airway score increased from 1.2±0.4 in saline lungs to 3.6±0.6, 2.5±0.3, and 3.3±0.5 in BLM lungs at 3, 6 and 22wk, respectively (**Figure 6K**, P=0.0059 by one-way ANOVA with non-parametric test; P=0.0073 for 3wk BLM lungs compared to saline lungs), alveolar score from 0.8±0.3 to 4.7±0.8, 4.8±1.1, and 4.6±0.8 (**Figure 6L**, P=0.0002 by one-way ANOVA with non-parametric test; P=0.0009, 0.0376 and 0.0261 for 3, 6 and 22wk BLM lungs compared to saline lungs, respectively), and vascular score from 1.9±0.3 to 2.8±0.2, 2.5±0.5, and 3.6±0.5 (**Figure 6M**, P=0.00318 by one-way ANOVA with non-parametric test; P=0.0393 for 22wk BLM lungs compared to saline lungs).

In addition to histological assessment, biochemical assessment of hydroxyproline content for collagen accumulation was performed to characterize our ferret model for preclinical applications according to guidelines of American Thoracic Society (24). A whole lobe of ferret lung was evaluated to be more presentative. As shown in **Supplemental Figure 2A**, although there was a trend of elevation (from 0.9±0.1 in saline and 3wk BLM lungs to 1.0±0.1 and 1.3±0.1 µg/mg lung in 6 and 22wk BLM lungs, respectively), hydroxyproline content was not significantly different along the time course (P=0.2856 by one-way ANOVA with non-parametric test). This is likely due to the patchy nature of the fibrotic area in the whole lung lobe as indicated by histological assessment, and potentially reduced amounts of free detectable hydroxyproline in the sample as fibrotic injury matured.

To visualize hydroxyproline distribution in the lung and assess this beyond free, detectable hydroxyproline, we applied immunohistochemistry with a hydroxyproline antibody that has been successfully utilized to follow hydroxyproline imprint in murine lungs (25). Representative images in **Supplemental Figure 2F-I** demonstrated greater presence of hydroxyproline in BLM compared to saline-exposed lungs at 22wk that was almost exclusively co-localized with fibrotic area as indicated by trichrome stain.

Furthermore, smooth muscle actin (SMA), was also detected by immunofluorescence. As shown in **Supplemental Figure 2J-M**, SMA was present in airways, vessels, and membrane of submucosal glands. Not only was it enhanced in interstitial lung post BLM exposure at both 3 (data not shown) and 22wk but also co-localized with fibrotic area as indicated by trichrome stain, highly consistent with hydroxyproline distribution.

### Aberrant Airway Remodeling in Areas of Fibrosis

Human data suggests aberrant metaplasia of ciliated cells in distal airway spaces could be an intermediate pathological phenotype in human IPF lungs (26, 27).

Progenitor basal-like cells originate from submucosal glands where *MUC5B* expression is natively high (28) and may be drivers of aberrant airway remodeling and distal “proximalization” in human IPF. Proximalization of the distal lung is a feature of human IPF that includes formation of honeycomb cysts, loss of club cells, and proximal airway markers (i.e. ciliated cells) in the distal lung indicative of aberrant regeneration (29–33), sometimes also referred to as ‘bronchialization’. Unlike mice, ferrets express MUC5B in a similar distribution to humans so we hypothesized that BLM ferrets may recapitulate mucociliary lesions if aberrant repair characterized by proximalization of distal spaces is key to sustained fibrosis phenotype (14). This could also be related to honeycomb cysts, pathologic structures lined with mucociliary, pseudostratified epithelium that express predominately MUC5B mucin (30) that rodent models have not consistently recapitulated.

It is hypothesized that distal airway metaplasia may be a precursor lesion to the honeycomb change; this could also be an early indicator of airway proximalization. To investigate this, we conducted dual IF for ciliated epithelium using the marker AT and non-ciliated bronchiolar epithelial cells using club cell secretory protein (CCSP). Airways double positive for CCSP^+^/AT^+^ would only be expected in select transitional regions of the tracheobronchial tree. An increase in the proportion of CCSP^+^/AT^+^ airways represent aberrant proximal reprogramming in the context of fibrotic injury. Our initial approach was a manual analysis in which we quantified the number of CCSP^+^/AT^+^ that met the threshold of >20% of airway cells staining positive for both CCSP and AT (examples in **Figure 7A,B** and **Supplemental Figure 1H,I,I’**). BLM-ferrets had increased proportion of CCSP+/AT+ airways compared to control (mean difference± SE: −33.11%±9.034; P=0.00219). This was accompanied by a decreased proportion of CCSP+/AT-airways compared to control (mean difference± SE: −25.5%±5.868; P=0.0015), and no meaningful change in CCSP-/AT+ airways (mean difference ± SE: −5.63%±5.575; P=0.113 (**Figure 7C**). There was no significant difference in the diameter of the CCSP+/AT+ airways between BLM (mean 171.7 µm ± 9.14) compared control (mean 205.5 µm ± 17.89; P=0.0955 by T-test (**Figure 7D)**.

**Figure 7.**
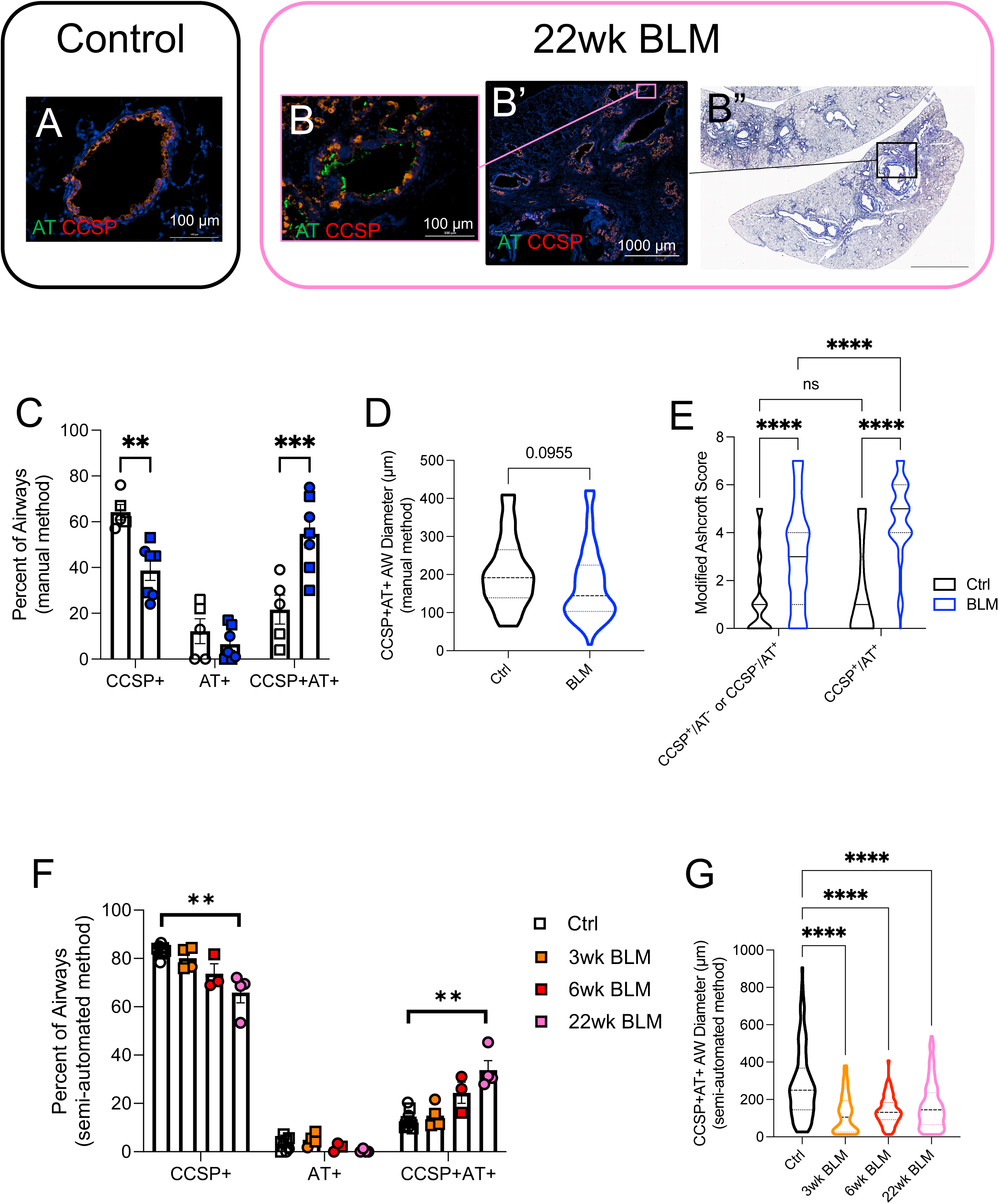
BLM-exposure induces aberrant airway remodeling in areas of fibrosis of ferret lungs. Ferrets were exposed to either saline or BLM and lung slices were obtained as in Figure 6. **A-B’’)** are representative images for ferret lungs 22wk post BLM exposure; immunofluorescence (IF) staining of alpha-tubulin (AT) and club cell secretory protein (CCSP) (A-B’) and Masson’s trichrome (B’’). Airway size and IF-staining signal for AT and CCSP were analyzed manually (C-E) and by semi-automated algorithm (F,G) as described in Methods. **C)** The proportion of airways stained CCSP+/AT-, CCSP-/AT+, and CCSP+/AT+ (n=156 airways analyzed across 5 control ferrets and n=222 BLM airways analyzed from 7 ferrets 3 or 6 weeks post BLM-injury using manual method). Airways were considered CCSP+/AT+ when threshold of >20% of cells was met for each stain. P=0.8701 for exposure and P<0.0001 for signal and interaction by 2way ANOVA with multiple comparisons (**P<0.01; ***P<0.001) by Šídák test. **D)** Size of double positive CCSP+/AT+ airways analyzed by manual method, P = 0.0955 by T-test, n = 27 control airways vs 104 BLM airways from airways analyzed in C). **E)** Modified Ashcroft score for airways analyzed in (C); P<0.0001 for exposure and IF-signal,P=0.0051 for interaction by 2way ANOVA with multiple comparisons (***P<0.001; ****P<0.0001) by Tukey’s test, n = 129 vs 114 and 27 vs 108 as control vs BLM for CCSP+/AT- or CCSP-/AT and CCSP+/AT+ groups, respectively. **F)** Percentage of airways electronically analyzed by algorithm described in methods for each ferret that are CCSP^+^, AT^+^, and CCSP^+^AT^+^. P=0.0011, 0.0625, and 0.0013 for CCSP^+^, AT^+^, and CCSP^+^AT^+^, respectively, by one-way ANOVA (Kruskal-Wallis test); N=8, 4, 3 and 4 for control, 3, 6 and 22wk post exposure, respectively; **P<0.01 by Dunn’s test. **G)** Size of double positive CCSP+/AT+ airways analyzed by algorithm described in methods. P<0.0001 (n=113, 80, 93 and 310 for control, 3, 6 and 22wk post exposure, respectively) by one-way ANOVA with multiple comparisons (****P<0.0001) by Dunnett test. n represents all IF+ airways irrespective of individual animal. Male ferrets represented by squares and female ferrets represented by circles.

We hypothesized that aberrant airway metaplasia would be the most abundant in areas with the most severe fibrosis, so we additionally examined the modified-Ashcroft score in BLM and control ferrets that approximate the CCSP^+^/AT^+^ airways as compared to airways without dual staining (**Figure 7E)**. Ashcroft score was significantly higher in CCSP^+^/AT^+^ airways than in single positive airways (CCSP^+^/AT^-^ or CCSP^-^/AT^+^) in the BLM cohort (mean [CI]; 4.6 [4.323 to 4.917] CCSP^+^/AT^+^ BLM vs 3.0 [2.649 to 3.439] single positive (CCSP^+^/AT^-^ or CCSP^-^/AT^+^) BLM; P<0.0001). Not surprisingly, both CCSP^+^/AT^+^ BLM and single positive (CCSP^+^/AT^-^ or CCSP^-^/AT^+^) BLM had significantly worse modified Ashcroft score than control airways. There was no difference between Ashcroft score in single positive (CCSP^+^/AT^-^ or CCSP^-^/AT^+^) and CCSP^+^/AT^+^ airways after control treatment (mean [CI]; 1.2 [0.926 to 1.492] single positive-control vs 1.5 [0.771 to 2.266] CCSP^+^/AT^+^ control, P=0.8474). To summarize, proximalization (as evidenced by an aberrant increase in proportion of CCSP^+^/AT^+^ airways) was increased following BLM exposure and associated with increased Ashcroft score in BLM-ferrets, suggesting that the fibrotic microenvironment influences epithelial cell fate and small airway repair processes.

To bolster the manual analysis and assess if proportion of double positive CCSP+/AT+ airways changes across the BLM time course, we developed a semi-automated quantitation method to not only detect airways containing IF signal but also measure their circularity and size as quality control metrics. As described in the supplemental methods, this quantitation includes five checkpoints that must be achieved. As a measure of quality control, circularity of detected airways was compared. As shown in **Supplemental Figure 3A,B**, IF+ airways presented similar distribution of circularity regardless of treatment, time point or stained protein. Total histological area analyzed was also similar between BLM and control conditions (**Supplemental Figure 3C**). This suggested that the quantitation method developed here consistently analyzed each lung slice without bias.

Applying this semi-automated method, the percentage of CCSP, AT, or MUC5B signal along the apical surface of the airway epithelia were estimated, therefore, the number of airways containing individual protein can be computed. Within all detected airways, **Figure 7F** demonstrates that the proportion of airways containing both CCSP and AT (CCSP+AT+) augments post BLM exposure (P = 0.0011 by one-way ANOVA). Although there were no significant difference between 3wk BLM (14.9±2.5, P>0.9999) vs saline (13.4±1.4%) or 6wk (24.4±4.4%, P=0.0988) vs saline, we observed an increased proportion of CCSP+AT+ at 22wk (33.8±3.9%, P = 0.0048 vs saline) post BLM exposure as fibrotic injury and airway remodeling ensued. As airways containing AT alone (AT+ only) maintained the same between treatment groups and along the time course, the number of airways containing CCSP alone (CCSP+ only) declined post BLM exposure, which consistently corresponded to the increase of CCSP+AT+ airways. As opposed to the manual analysis which examined a fewer number of airways (i.e. Figure 7D), we found that the average size of CCSP+AT+ airways was smaller post BLM exposure (**Figure 7G**, mean±SEM from 280±18 for control to 123±11, 139±8 and 168±7 µm for 3, 6 and 22wk post BLM exposure, respectively, P<0.0001 by one-way ANOVA), suggesting the proximalization phenotype was occurring in distal airway spaces.

Beyond analyzing individual airways, we also evaluated as a mean value for each individual animal controlled for area examined. The quantitative number of airways that contain CCSP increased from 28±3 baseline to 68±21 (P=0.0081) at 22wk post BLM exposure (**Supplemental Figure 4A**); AT also increased from 4±1 baseline to 22±5/cm^2^ (P=0.0040; **Supplemental Figure 4B**). Similarly, MUC5B had a trend towards increased expression by animal at 22 wks (15±3 BLM vs 4±1/cm^2^ control, P=0.0571; **Supplemental Figure 4C**). Overall, this demonstrated distal airway metaplasia representative of distal airway proximalization that worsened over time in BLM-exposed ferrets.

Humans with IPF have MUC5B-rich honeycomb cysts; BLM exposed ferrets also exhibited this abnormality (**Figure 6A-H, Supplemental Figure 1A-D**). MUC5B^+^ AT^+^ airways 22 wks after BLM exposure had a reduced airway diameter compared to controls (**Supplemental Figure 1G**; 228±12 vs saline 394±28µm, P<0.0001 by T-test).

In addition to analysis of defined airways, BLM exposure also caused CCSP+ gland-like structures in proximity to the alveolar space that were intentionally excluded by the automated analysis (**Figure 7B, B’**). Additionally, IF staining revealed increased CCSP+ (but not MUC5B+) submucosal glands following BLM exposure compared to saline (**Figure7B’**). These structures may reflect genesis of cystic metaplasia seen in IPF and BLM exposed ferrets (**Supplemental Figure 1C,D**).

## Discussion

IPF is a fatal lung disease characterized by restrictive physiology and progressive respiratory failure as the functional alveolar-capillary units are destroyed by deposition of extracellular matrix in the interstitium. The prevalence of IPF is increasing and it affects nearly 0.5% of people over 65 in the United States (34–36). Though the disease is idiopathic, the most dominant risk factor for developing IPF is the promoter variant rs35705950 for MUC5B (2, 37). Patients with IPF demonstrate excess MUC5B mucin in their respiratory bronchioles and have MUC5B filled honeycomb lesions (30). However, it is unknown how mucin microenvironments contribute to pathogenesis and dysregulated repair in IPF. Using ferrets, a species that exhibits native expression of the MUC5B promoter and tissue and cell type distribution of MUC5B expression that resembles humans, we establish a novel ferret model of pulmonary fibrosis that exhibits physiologic, radiographic, and histological evidence of persistent disease and clear evidence of aberrant remodeling of the distal airways, clear indicators of human disease absent in rodents and strong affirmation of the importance of MUC5B expression in human IPF.

A substantial obstacle for developing appropriate animal models of respiratory disease relates to the genetic, anatomic, and functional variations among species. Notably, submucosal gland expression in rodents is limited to the proximal trachea. At the same time, in humans they are present in major airways and bronchioles, which has significant implications for the ability of rodents to model human diseases with mucus-related pathophysiology. For example, murine models of cystic fibrosis accurately replicate the ion-transport abnormalities and disease phenotype in the gastrointestinal tract (38), yet alterations in airway mucus transport that are so devastating in humans remain largely absent. Similarly, mouse models of COPD do not recapitulate problems of mucus retention and bronchitis (39). Recently, ferret models of both CF and COPD-bronchitis have been shown to develop human-like lung disease (16, 17, 40). The COPD-bronchitis ferret model that we developed previously exhibits cough and airway obstruction as well as pathologic evidence of chronic mucus hypersecretion (17). In addition, CF ferrets exhibit CFTR-dependent secretion in trachea and a predisposition to lung infections during the early postnatal period (40–42). These models demonstrate how lung anatomy and lung cell biology of ferrets provide better models of pulmonary disease given their similarity to humans. The development of the pulmonary fibrosis ferret model provides the first opportunity to define respiratory mechanics and repair mechanisms in a model that natively expresses the IPF risk conferring promoter variant rs35705950 for MUC5B.

In this study, we describe a novel BLM-induced ferret model of pulmonary fibrosis and describe a persistent phenotype that recapitulates hallmarks of human IPF through 22wks with a single installation of BLM. BLM was first used experimentally to establish pulmonary fibrosis in mice in 1974 (44) and the BLM-mouse is now the best characterized animal model of pulmonary fibrosis (12). A recent Workshop report by the American Thoracic Society recommended a single-dose of BLM-mouse administered via the oropharynx for preclinical studies (45). We sought to establish a ferret model that could be comparable to the existing literature, thus, we decided to install a single dose of BLM I.T., in a weight and length specific manner (to control for sexual dimorphism of ferrets). We conducted dose-response studies (data not shown) informed by the murine literature and selected 5U/kg as the optimal dose. Of note, 50% mortality has been reported in mice treated with 5U/kg BLM (46), but did not cause mortality in our ferret model.

A major strength of the ferret model is the ability to incorporate longitudinal pulmonary function measurements and serial µCT imaging to track disease progression throughout the experiment that are highly relevant to humans, in addition to histological and biochemical outcome measures in preclinical models. The BLM ferret model demonstrated restrictive lung physiology that was sustained through 22-weeks as evidenced by downward and right shifted PV loops with significantly decreased slope of the deflation limb, sustained decrease in inspiratory capacity, increased whole lung elastance, and decreased compliance. In total, the pathophysiology we report is concordant with the sustained fibrotic remodeling we observed histologically and biochemically. Additionally, the restrictive change occurred simultaneously with a decrease in pulse oxygen saturation.

Radiodensity of the lungs, as quantified with volumetric µCT analyses, revealed a significant, sustained increase in opacification by P85 through 22 weeks compared to baseline in the BLM-ferrets. Volumetric CT analysis of human IPF scans have been shown to closely correlate with lung physiological parameters and is useful in detecting composite unit stage (53). Future directions include stratifying individual ferrets to see if there are different persistent vs progressive fibrosis phenotypes based on their functional and imaging data, taking advantage of the fact that ferrets are an outbred species.

In concert with persistent physiologic abnormality, we observed clear evidence of pulmonary fibrosis as evident by general histology and that matured over time within the ferret lung, and eventually generated pathologic hallmarks of human IPF, such as MUB5B-enriched cystic structures that resemble honeycomb change. Importantly, we also identified key features of aberrant repair and airway remodeling evident in human IPF that have not been reported in murine models (12, 45). This was accompanied by elevation of hydroxyproline, an important marker of collagen content in preclinical models (47), but lacks spatial information. Although free hydroxyproline levels in whole lung lobes reverted after 12 weeks, histological detection of hydroxyproline indicated persistent abnormally high expression of hydroxyproline throughout the injured lung; we suspect reduced detection in lung homogenate may be related to sampling error, geographic disease heterogeneity, and potentially reduced detection of free hydroopxyproline as the disease matures.

Recently, Verleden and colleagues found that the loss of terminal bronchioles was associated with the appearance of fibroblastic foci and an increase in the volume fraction of fibroblastic foci in their sophisticated microCT and histological analysis of IPF explants (32). In ferrets, we observed metaplastic ciliation surrounding fibroblastic foci and proximalization in areas with more severe fibrosis as evidenced by increased co-expression of CCSP and AT in distal airway spaces that has been defined previously (33, 54, 55). We also observed the formation of honeycomb cyst and fibrotic foci, pathologic features seminal to human IPF (29, 30, 56, 57) and compatible with the aberrant remodeling of the airways. These observations supports an emerging paradigm shift that proximalization is not simply a consequence of fibrosis, but may be a driver and play a role in the sustained disease we observed in our model (32, 33). The model is well suited to investigate the time course for the loss of normal proximal-peripheral differentiation of pulmonary epithelial cells with implementation of molecular techniques beyond the scope of the present work.

The increased proportion of CCSP+ airways in BLM-exposed ferret lungs is an interesting phenotype and may allow future mechanistic elucidation of the role of club cells in fibrosis pathogenesis; work by Fukomoto and colleagues demonstrated phenoconverted club cells which may participate in the initiation and progression of fibrosis via bolstered proliferative and migratory ability (31) and Reynaud and colleagues characterize the bronchiolization from human ILD lung biopsies by loss of SCGB1A1(+) bronchiolar-specific club cell density (33). Caveats from these works include the small sample size of patients and limited biopsy material which was predominantly small airways.

Though the fibrosis in the ferret model was dependent on deposition of BLM and therefore most intense around the proximal airways, we overcome some limitations of the human studies such that whole lobes, including all airway sizes, were included in analysis. Other limitations of our IF quantitation method were described in the Supplemental Material. Bleomycin exposure is the known, experimentally controlled nidus for fibrotic injury in our model system, and this is of course distinct from the complex interplay of genetic, environmental and idiopathic factors that lead to development cause in human IPF. However, the shared mechanisms for pathologic remodeling such as proximalization and honeycomb cyst formation, can be elucidated leveraging a ferret model that recapitulates hallmarks of human disease in combination with advanced staining and quantitation methodology. We noted more severe restriction in female ferrets than males. We do not attribute the difference to sex hormones given that the jills were spayed and the hobs were neutered. Instead, we suspect the more severe female phenotype to be related to airway diameter and consequent BLM deposition. Though weight and length specific dosing were employed, the female airways’ smaller caliber could have influenced dose delivery. Elucidating mechanism underlying these sex differences is of interest in the future. Future studies are poised to elucidate the role of MUC5B in pulmonary fibrosis pathogenesis with mechanistic studies that alter the mucin microenvironment pharmacologically and could more definitively establish MUC5B as a therapeutic target in the disease.

In summary, we established and characterized a novel ferret model of pulmonary fibrosis which demonstrates sustained fibrosis and evidence of airway remodeling that featured airway bronchiolization of the distal airway spaces, and clear resemblance of human airway morphology present in IPF. The BLM-ferret model represents an important advance that can help advance preclinical animal models of pulmonary fibrosis.

## Acknowledgements

We sincerely thank UAB Pathology Core Research Laboratory for IHC and IF staining and UAB Comparative Pathology Laboratory for tissue processing, slicing, and H&E and trichrome staining. We also thank UAB High Resolution Imaging Facility and the imaging core of UAB Cystic Fibrosis Research Center for microscopy and µCT imaging. We acknowledge useful contributions from the NHLBI U01 IPF Consortium.

## Supplemental Figure Legends

**Supplemental Figure 1:**
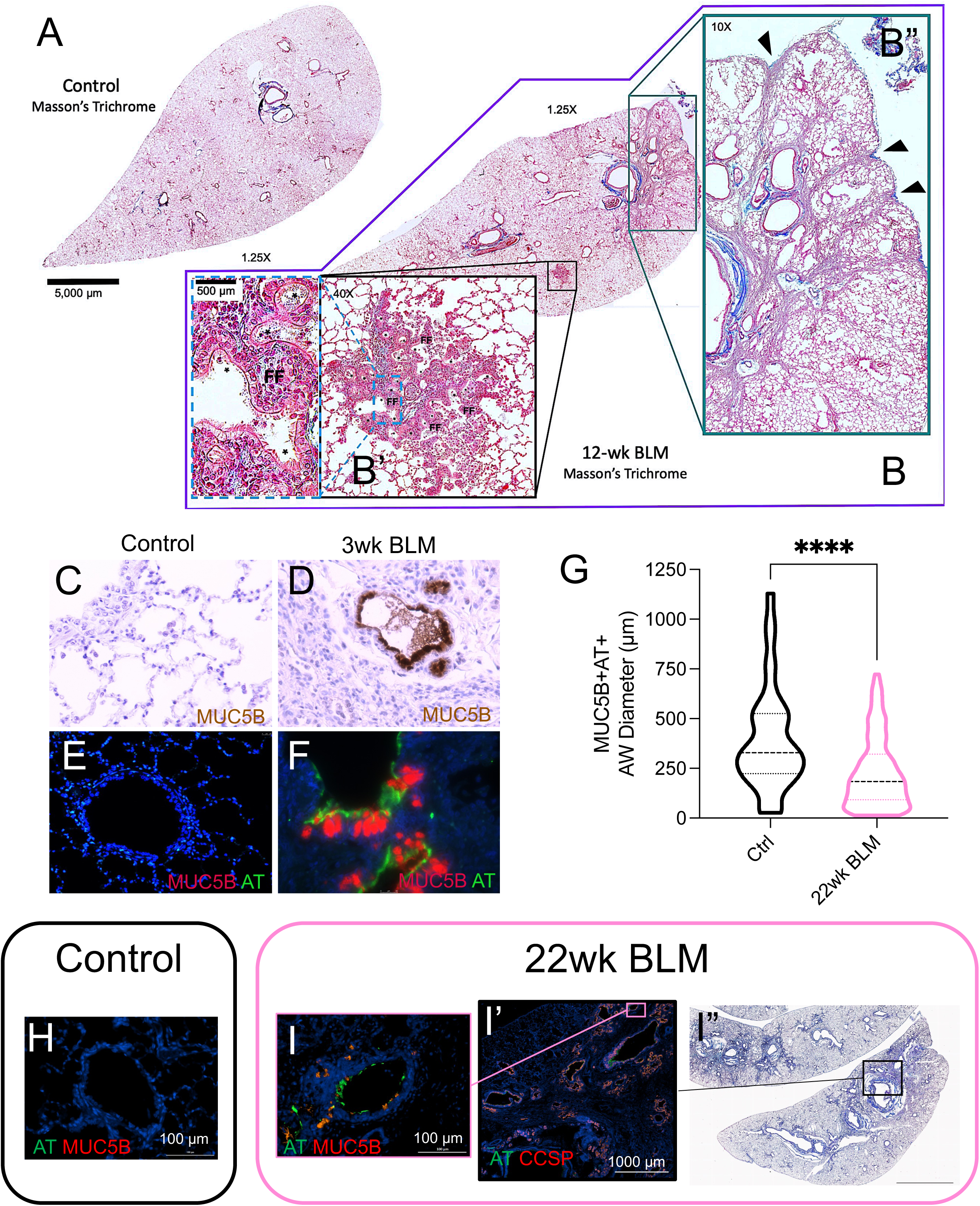
BLM ferret recapitulates hallmarks of IPF including fibroblastic foci, MUC5B+AT+ mucociliary cystic spaces, and proximalization of small airways. Ferrets were exposed to saline or BLM and lung slices were obtained as in Figure 6. Masson’s trichrome (**A-B’’, I’’**), immunohistochemistry (IHC) of mucin 5B (MUC5B; **C,D**), immunofluorescence (IF) of alpha-tubulin (AT) and MUC5B (**E-I’**) were stained, IF+ airways and their diameters were analyzed by semi-automated algorithm (**G**) as described in Methods. **B’)** Black inset and dotted blue inset showed fibrotic foci (FF) at high power and * denote areas with abnormal ciliation. **B”)** Green inset and arrowheads demonstrated retraction at the visceral pleural surface with interstitial fibrosis extending into the parenchyma. Airway centric fibrosis is apparent. Immunohistochemical stain for MUC5B (brown) with hematoxylin counterstain in **C)** saline control with normal alveoli and interstitium and **D)** BLM treatment with MUC5B-rich honeycomb cysts. Immunofluorescence stains for MUC5B (red) and AT (green) in **E)** saline control and **F)** BLM treatment. **G)** Semi-automated algorithm analysis of ferret airways 22wk post exposure, of which representative images shown in **H-I’’).** P<0.0001 (n=83 and 225 for control and BLM, respectively) by T-test. n here represents all IF+ airways irrespective of individual animal.

**Supplemental Figure 2:**
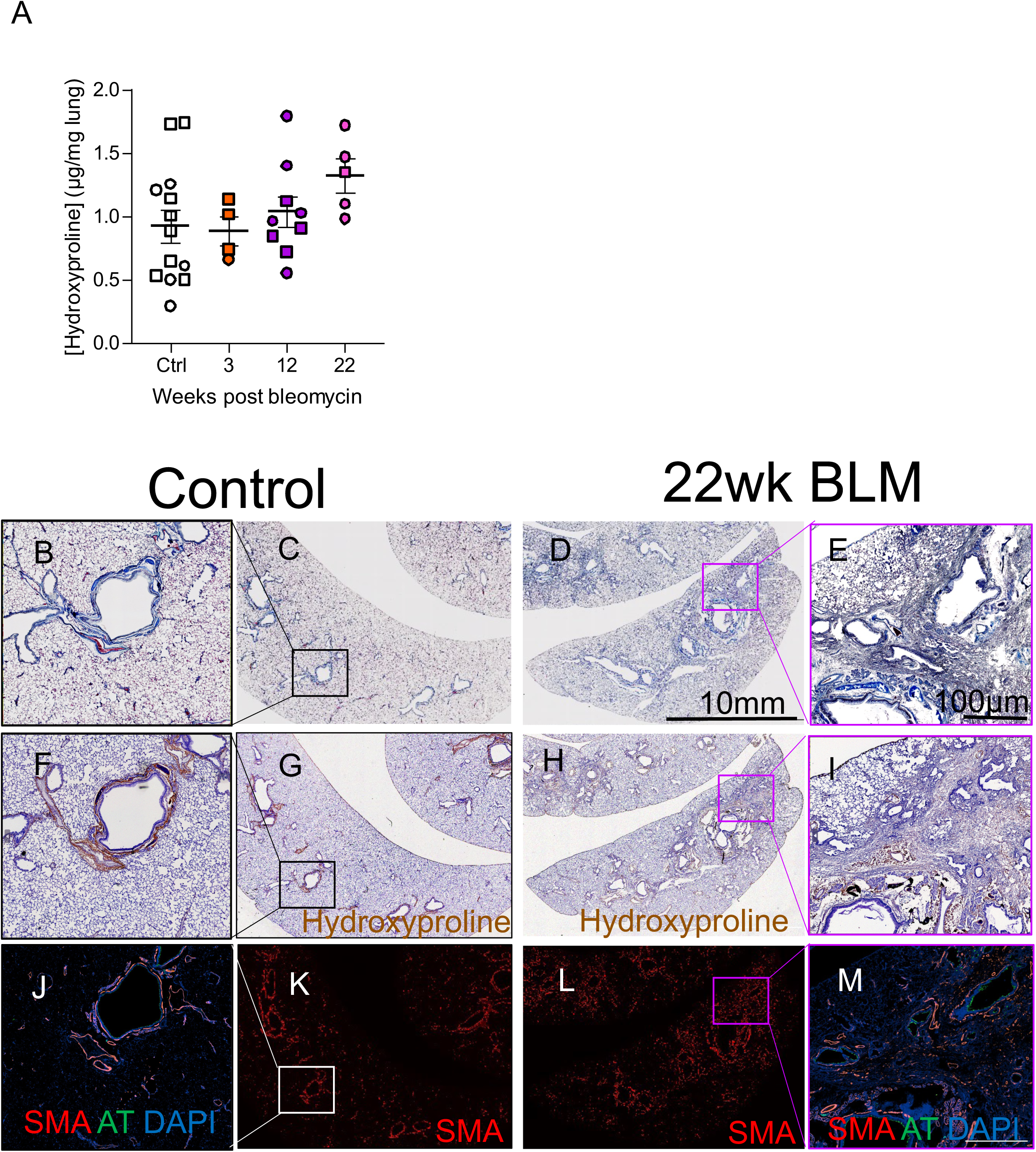
Biochemical and histologic evaluation of fibrotic injury. Ferrets were exposed to saline or 5U/kg BLM as in Figure 1, one lobe of their right lungs were collected at 3, 12 and 22wk and processed for hydroxyproline assay (**A**), and one of the lower lobes at 22wk BLM compared to control for Masson’s trichrome (**B-E**), hydroxyproline IHC (**F-I**), and IF of alpha-smooth muscle actin (SMA) and alpha-tubulin (AT; **J-M**) as described in Methods. P=0.2856 by one-way ANOVA (Kruskal-Wallis test); N = 13, 4, 9 and 5 for saline, 3, 12 and 22wk group, respectively (A). Squares indicate males and circles females.

**Supplemental Figure 3.**
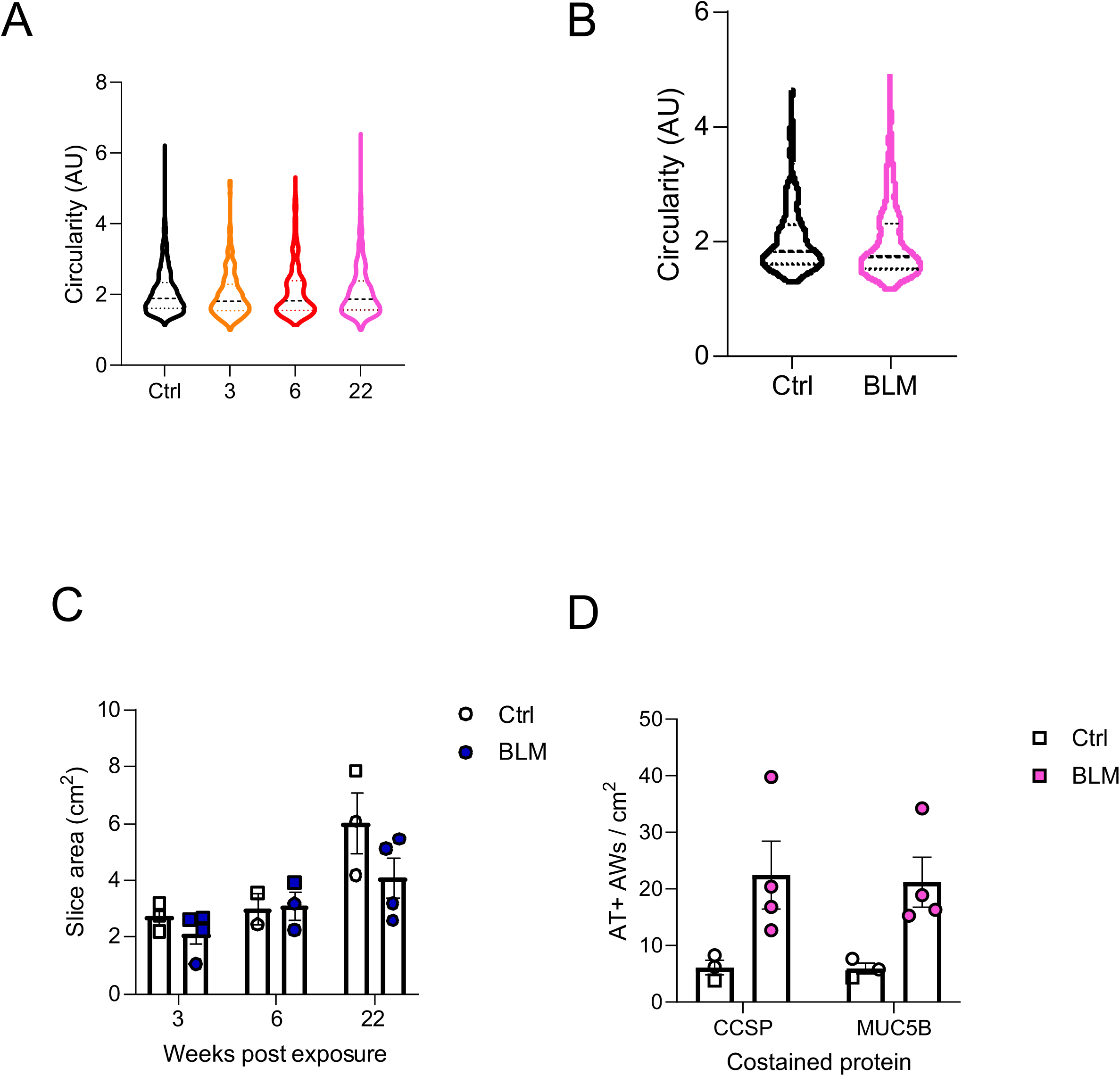
Assessments of the quality control for histology and IF methodology. Saline or BLM exposed ferret lung slices were obtained and stained as in Figure 7. IF+ airways in CCSP/AT (**A, C**) and MUC5B/AT (at 22wk post exposure; **B and D**) co-stained lung slices were selected and their circularity were calculated as described in Methods. P = 0.1994 (n=918, 486, 363 and 955 for control, 3, 6 and 22wk post exposure) by one-way ANOVA (A), and 0.6252 (n=105 vs 311 for control vs BLM at 22wk post exposure, respectively) by T-test (B). n here represents IF+ airways irrespective of individual animal. The surface area of lung slices used for CCSP and AT costain by IF were estimated as described in Methods. C) Total surface area examined by time point and exposure group. D) Comparison of total AT+ AWs counted by semi-automated quantitation method from two different datasets defined by co-stain method. N=3 vs 4, 2 vs 3 and 3 vs 4 for saline vs BLM at 3, 6 and 22wk post exposure, respectively.

**Supplemental Figure 4.**
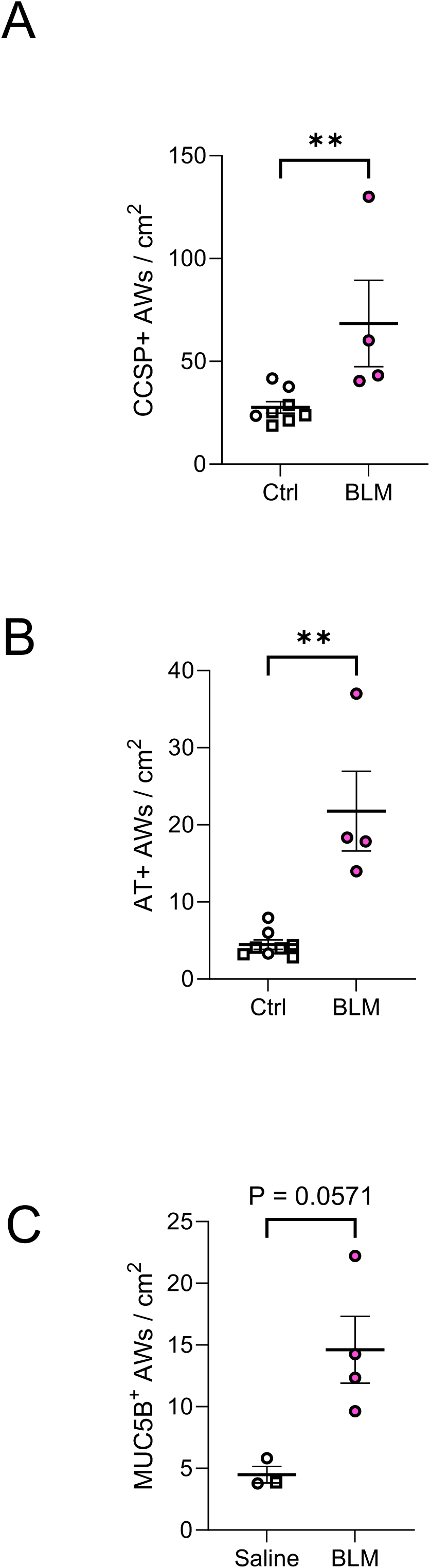
Number of IF+ airways in ferret lungs post saline or BLM exposure. Saline or BLM were exposed, lung slices were obtained and stained, and airways that were CCSP+ (**A**), AT+ (**B**), and MUC5B+ (**C**) at 22wk post BLM or control were counted as in Figure 6. Number of airways were normalized to the lung slice area evaluated for each animal. P = 0.0081, 0.0040 and 0.0571 in A, B, and C, respectively, by Mann-Whitney test. Squares indicate males and circles females.

## Supplemental Methods

### Lung slice preparation

Longitudinal-cut slices at up to four locations evenly distributed from the proximal to distal airway to best represent the whole fixed lung lobe were stained for H&E, Masson’s trichrome, hydroxyproline immunohistochemistry (IHC) and immunofluorescence. Total 2.7±0.3 vs 2.1±0.4, 3.0±0.6 vs 3.1±0.5 cm^2^ of lung slice per animal was evaluated for ferrets 3 and 6 wk post exposure (saline vs BLM), respectively, while larger pieces 6.0±1.1 vs 4.1±0.7 cm^2^ (saline vs BLM) for 22wk to ensure persistency of disease features. At each time points, there was no difference on evaluated area of lung slices between saline and BLM groups (**Suppl. Figure 3C** for CCSP/AT IF lungs as an example).

### Histological Analysis

Histology was analyzed by a board-certified veterinary anatomic pathologist blinded to experimental groups. Masson’s trichrome stained lung slices were evaluated for both fibrotic area (% of the whole slices) and Ashcroft score of up to five representative 20X areas using standardized method (1). Other lesions particularly inflammation in H&E stained lung slices were semi-quantitated for airway, alveolar, and vascular sublocations following published criteria and slightly modified algorithm (2).

Briefly, the total score for each sublocation (0-11, 0-12, and 0-8 for airway, alveolar, and vascular score, respectively) was the sum of the extent of affected area (0 – 4 with 0 = none, 1 < 10%, 2 = 10 – 25%, 3 = 25 – 50%, and 4 > 50% of lung surface area displaying pathologic lesion spectra, respectively), severity according to defined criteria (0 – 4 for airway and alveolar and 0 – 3 for vascular sublocation), and location-specific criteria including bronchial epithelial hyperplasia (0–3) for airway score, type II pneumocyte hyperplasia (0–3) and edema/protein/hemorrhage/fibrin (present = 1 and absent = 0) for alveolar score, and necrotizing vasculitis/thrombi (present = 1 and absent = 0) for vascular score, respectively.

### Biochemical assay and immunohistochemistry of hydroxyproline

Flash-frozen lung lobes were grinded, and the powder were mixed under liquid nitrogen. 0.1g/L of lung powder in water was homogenized by PowerGen 125 (Fisher Scientific). Hydroxyproline content was measured by hydroxyproline assay kit (cat. # MAK008 from Sigma) following manufactural instructions.

Unstained lung slices were labeled with hydroxyproline by UAB Pathology Core Research Laboratory. Standard IHC process was applied with antigen retrieval by 0.01 M Tris buffer containing 1mM EDTA (pH 9), 1:200 dilution of anti-hydroxyproline antibody (cat. # 73812 from Cell Signaling Technology) (3) and 1:1000 dilution of goat anti-rabbit IgG H&L secondary antibody conjugated with HRP (Abcam cat. # ab6721).

### Immunofluorescent Staining

Unstained lung slices were labeled with αcetylated-tubulin (AT), club cell secretory protein (CCSP), mucin 5B (MUC5B) and smooth muscle actin (SMA) by UAB Pathology Core Research Laboratory. Standard immunofluorescence (IF) process was applied with antigen retrieval by 0.01 M Tris buffer containing 1mM EDTA (pH 9), blocking by 5% normal donkey serum (cat. # D9663 from Sigma), 1:50 dilution of primary antibodies including AT (cat. # T7451 from Sigma-Aldrich), CCSP (cat. # 10490-1-AP from Proteintech), MUC5B (cat. # HPA008246 from Sigma), SMA (cat. # ab5694 from Abcam), 1:100 dilution of secondary antibodies including donkey anti-mouse IgG Northern Lights™ NL493-conjugated antibody (cat. # NL009 from Fisher) and donkey anti-rabbit IgG NorthernLights™ NL557-conjugated antibody (NL004 from Fisher). The slides were covered with cover-glasses using ProLong™ Gold Antifade Mountant containing DAPI (cat. # P36931 from Fisher).

As a quality control measure, CCSP/AT/DAPI-stained serial sections from two saline and two BLM-exposed animals were compared, demonstrating reproducible, consistent localization between the technical replicates.

### Semi-Automated Analysis of Immunofluorescence

The percentage of AT, CCSP or MUC5B signal along the apical surface of the airway epithelia (AT%, CCSP% or MUC5B%) were assessed using a comprehensive methodology involving image processing in Image J (NIH, Bethesda, DC) and lab-made software using LabVIEW™ (National Instrument, Austin TX).

For the dual-staining of AT and CCSP, 4X stitched raw images of individual DAPI, CCSP or AT staining were processed in Image J to extract signal above a defined threshold and generate individual masks. Three masks corresponding to DAPI, CCSP, and AT signals for each slice were overlaid, and the CCSP+ or AT+ airways were manually selected based on comparison with the original IF images. This selection process generated airway masks containing exclusively CCSP+ or AT+ airways, with manual correction applied to rectify any false shapes caused by debris or misidentified alveolar spaces observed in the original IF images. The corrected airway masks were used to calculate various parameters (number, circularity, lumen perimeter, and lumen area) of the airways utilizing LabVIEW scripts. Heywood circularity, defined as the perimeter divided by the circumference of a circle with an equivalent area (4), was employed for circularity measurements. The cross-sectional perimeter of airways was estimated as the perimeter of the largest circle that fits within the airway lumen in the airway mask, providing an indication of airway size in terms of diameter.

To compute the AT% (percentage of AT signal along airway apical surface), the corrected airway mask was overlaid with the AT mask. For CCSP%, considering that club cells (CCSP+) are located outside the airway lumen, the corrected airway mask was dilated three times (airway mask 3X) to encompass majority of the CCSP signal within the masks of each airways. Airway mask 3X was manually corrected if merging of airways occurred due to dilation, and was confirmed to contain the same number of airways as the non-dilated airway mask. CCSP% was calculated from the overlay of the airway mask 3X and the CCSP mask using LabVIEW scripts.

Two threshold settings (low as mean + 4-5SD and high as mean + 6-8SD) for generating AT mask and three (low as mean + 5-8SD, high as mean + 3-5SD and auto local) for CCSP were applied. The final AT/CCSP% applied in the analysis was manually selected from these results with different threshold settings or adjusted to correct false reads by comparing them to the original IF images.

The same analysis procedures were applied to AT and MUC5B dual-staining with thresholds of mean + 5-7SD and mean + 7-10SD for generating AT and MUC5B mask, respectively.

Serial cut of the same paraffin-embedded blocks/lung lobes (22wk) were stained for either CCSP or MUC5B in combination with AT. As expected, total AT+ AWs counted by our quantitation method for the cut from the same lung lobe (saline or BLM) were the same when either costained with CCSP or MUC5B (**Suppl. Figure 3D**; P=0.8765 by 2-way ANOVA). This demonstrated the consistency of our staining process and quantitation methodology.

## Supplementary Discussion

General caveats of histology and IF staining using tissue slices such as (but not limited to) their representability of the whole lung, the tissue integrity by cutting and placing onto slides, and occasionally ambiguous cut of airways, apply here. For IF quantitation method, there are important caveats. First, using cross-sectional perimeter/diameter to represent airway size avoids overestimation using lumen perimeter/diameter for majority of airways where the cut was between cross-sectional and longitudinal (the more longitudinal the cut is, the greater overestimation of the actual airway size is). However, this technique has the potential to underestimate airway size for small amount of airways that were cut cross-sectionally. Under-inflation to a minor degree was observed in rare cases but was not systematic between conditions.

Second, it is possible that some small airways were neglected during airway selection by Image J. This amount is negligible except in the case with massive increase of CCSP^+^ cells, which would bias results against a finding. Lastly, the current method has the potential to not distinguish clusters of CCSP^+^ areas that are proximal to the alveoli in transitional small airways; all sections for automated analysis were also manually examined to assure this was handled in an unbiased fashion, whereas larger CCSP^+^ regions that were not clearly airways were reported separately.

